# Mapping the structural coverage of *Arabidopsis thaliana* plant developmental proteins: Insights from Experimental and AlphaFold Approaches

**DOI:** 10.64898/2026.05.31.729038

**Authors:** Surabhi Shriram Rode, Karthik Sudarsanam, Hemal Bhalla, Ashutosh Srivastava, Subramanian Sankaranarayanan

## Abstract

**Background:** Plant development is a multifaceted process governed by intricate protein regulatory networks. High-throughput sequencing methods have vastly expanded plant transcriptomic and proteomic datasets, yet there is a large discrepancy between structural information for plant developmental proteins and the UniProt sequence entries. Advances in X-ray crystallography, NMR spectroscopy, and Cryo-EM have enabled the determination of protein complex structures and their dynamics. AI-driven tools like AlphaFold have revolutionized analysis of protein structural intricacies. However, available three-dimensional structural models predominantly prioritize the human proteome and other mammals over plants. Assessing structural coverage of plant developmental proteins is thus essential to identify research gaps, guide structure-function studies, and advance agriculture.

**Results:** Here, we focus on mapping the structural coverage of developmental proteins in *Arabidopsis thaliana*. We observed a substantial disparity in the Protein Data Bank (PDB) representation of *Arabidopsis thaliana* proteins compared to those of *Homo sapiens.* Our analysis identified 16,389 reviewed UniProt entries, of which only 1,038 have experimentally determined structures. Functional mapping using PlantGSEA revealed 3,485 proteins associated with plant developmental processes; of which only 337 (9.67%) have experimentally determined structures. In contrast, analysis of the AlphaFold database showed that 69.85% of the 39,278 *Arabidopsis thaliana* UniProt protein entries have predicted structures. Notably, all 3,485 plant developmental proteins (100%) from *Arabidopsis thaliana* are covered by AlphaFold models. The substantially higher structural coverage provided by AlphaFold for *Arabidopsis thaliana,* relative to *Homo sapiens,* highlights the strength of computational approaches in addressing the challenges of structural studies of difficult-to-crystallize proteins. Furthermore, 79.15% of reviewed *A. thaliana* protein models exhibit high confidence (pLDDT > 70), indicating reliable structural predictions. Although the experimental structural coverage of *Arabidopsis thaliana* developmental proteins remains limited, AlphaFold has markedly expanded the accessible structural landscape.

**Conclusion:** This study investigated the structural coverage of *Arabidopsis thaliana* plant developmental proteins, underscoring the critical need for structural studies using both experimental and AlphaFold approaches. It provides research directions for bridging the knowledge gap in understanding molecular mechanisms of plant development.

## Background

Plant growth and reproduction are complex processes regulated by a dynamic interplay of genetic and environmental factors. The plant life cycle is continuous and comprises a series of sequential events, including seed germination, plant growth, maturation, reproduction, and senescence [1]. Following reproduction, seeds are formed and subsequently germinate, initiating a new developmental cycle that progresses through growth and maturation before returning to reproduction and senescence. These developmental processes are regulated by a complex interplay of signaling molecules, proteinaceous receptors, and various biochemical pathways, responding to intrinsic and extrinsic factors [2]. Understanding these mechanisms requires detailed investigation of the proteins and protein complexes that govern them. Such knowledge has significant implications for agriculture, enabling the development of crops with higher yields, improved quality, and enhanced resistance to biotic and abiotic stresses. Structural characterization of proteins, including their post-translational modifications (PTMs) and interaction networks, provides critical insights into their molecular functions, revealing how proteins interact with DNA, other proteins, and cellular components.

Advances in protein crystallography, beginning with the determination of myoglobin’s structure in 1958 [3], have enabled the resolution of numerous protein structures from diverse organisms, including animals, microbes, and plants. The first crystal structure of a plant protein, papain, was reported in 1968 by J. Drenth et al., offering important insights into its catalytic mechanism [4]. Despite the wide diversity and functional importance, plant proteins remain underrepresented in structural databases compared to mammalian proteins [5]. Recent developments in structural biology techniques, including nuclear magnetic resonance (NMR) spectroscopy and cryo-electron microscopy (cryo-EM), and also bioinformatic tools have accelerated protein structure determination. In particular, cryo-electron microscopy (Cryo-EM) has transformed the study of large protein complexes, enabling high-resolution visualization of their architecture and interactions [6]. However, structural characterization of plant proteins remains challenging due to their complexity, multimeric organization, extensive PTMs, and the prevalence of membrane-associated receptors, which complicate experimental analyses [7]. The emergence of artificial intelligence–driven approaches, particularly AlphaFold and its subsequent versions, has revolutionized protein structure prediction since 2018. These deep learning models have significantly improved the accuracy and speed of structural predictions, providing reliable models for proteins that are difficult to resolve experimentally (Figure 1) [8, 9]. This transition from the pre- to post-AlphaFold era has markedly expanded structural coverage across proteomes, including those of plant developmental proteins (Figure 1).

**Figure 1:**
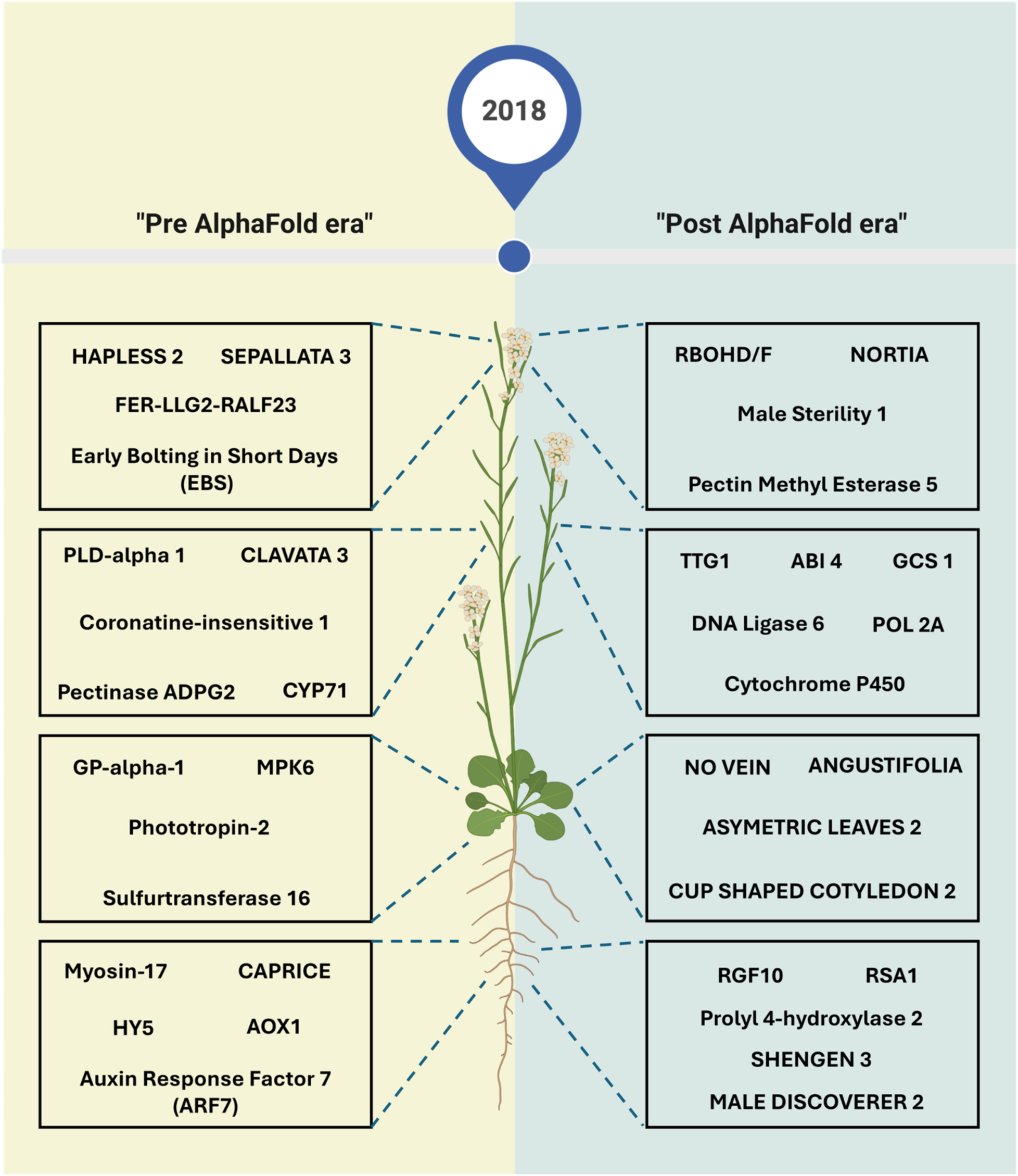
Overview of the effect of AI-driven AlphaFold discovery on the structural coverage of various proteins involved in plant development.

Nevertheless, structural data in the Protein Data Bank (PDB) remain disproportionately skewed toward the human proteome, highlighting a critical gap in plant protein research. Addressing this imbalance is essential for advancing our understanding of plant physiology and for enabling structure-guided agricultural innovations.

Here, we have examined the structural coverage of proteins involved in plant developmental processes, with a particular focus on *Arabidopsis thaliana*, a widely used model organism. It identifies key gaps in current structural knowledge and emphasizes the importance of integrating structural and functional studies. Such efforts will be instrumental in driving biotechnological advancements aimed at developing improved and resilient crop varieties.

### Current status of structural studies on proteins involved in plant development

Recent advancements in high-throughput sequencing technologies, multi-omics approaches, and bioinformatics tools have substantially expanded the availability of transcriptomics and proteomics data [10]. These developments have enhanced our ability to explore protein sequence diversity, assess sequence similarity to known proteins, and classify proteins, thereby providing valuable insights into their biological functions. However, the classic “protein folding problem”—where primary amino acid sequence alone does not fully determine biological function—underscores the need to investigate protein three-dimensional structures and functional domains. X-ray crystallography remains the gold-standard for protein structure determination and has driven major breakthroughs in structural biology [11]. Complementary techniques such as nuclear magnetic resonance (NMR) spectroscopy enable the study of protein structures and conformational dynamics in solution, offering insights into molecular interactions essential for biological function [12]. More recently, cryo-electron microscopy (cryo-EM) has overcome several limitations of traditional methods by eliminating the need for crystallization and simplifying sample preparation compared to NMR approaches requiring isotopic labeling [13]. Cryo-EM has proven particularly powerful for resolving large protein complexes and membrane proteins, which have historically been difficult to characterize [14].

The number of experimentally determined protein structures has steadily increased, with the Protein Data Bank (PDB) now containing approximately 245,715 entries (Figure 2a), reflecting significant advances in techniques such as X-ray crystallography, cryo-EM, and NMR spectroscopy. Despite this progress, plant proteins remain underrepresented. Only 2,265 (0.92%) PDB entries correspond to *Arabidopsis thaliana*, a key model organism, whereas 71,186 (28.97%) are derived from *Homo sapiens* (Figure 2b). This disparity highlights a significant gap in plant structural biology.

**Figure 2:**
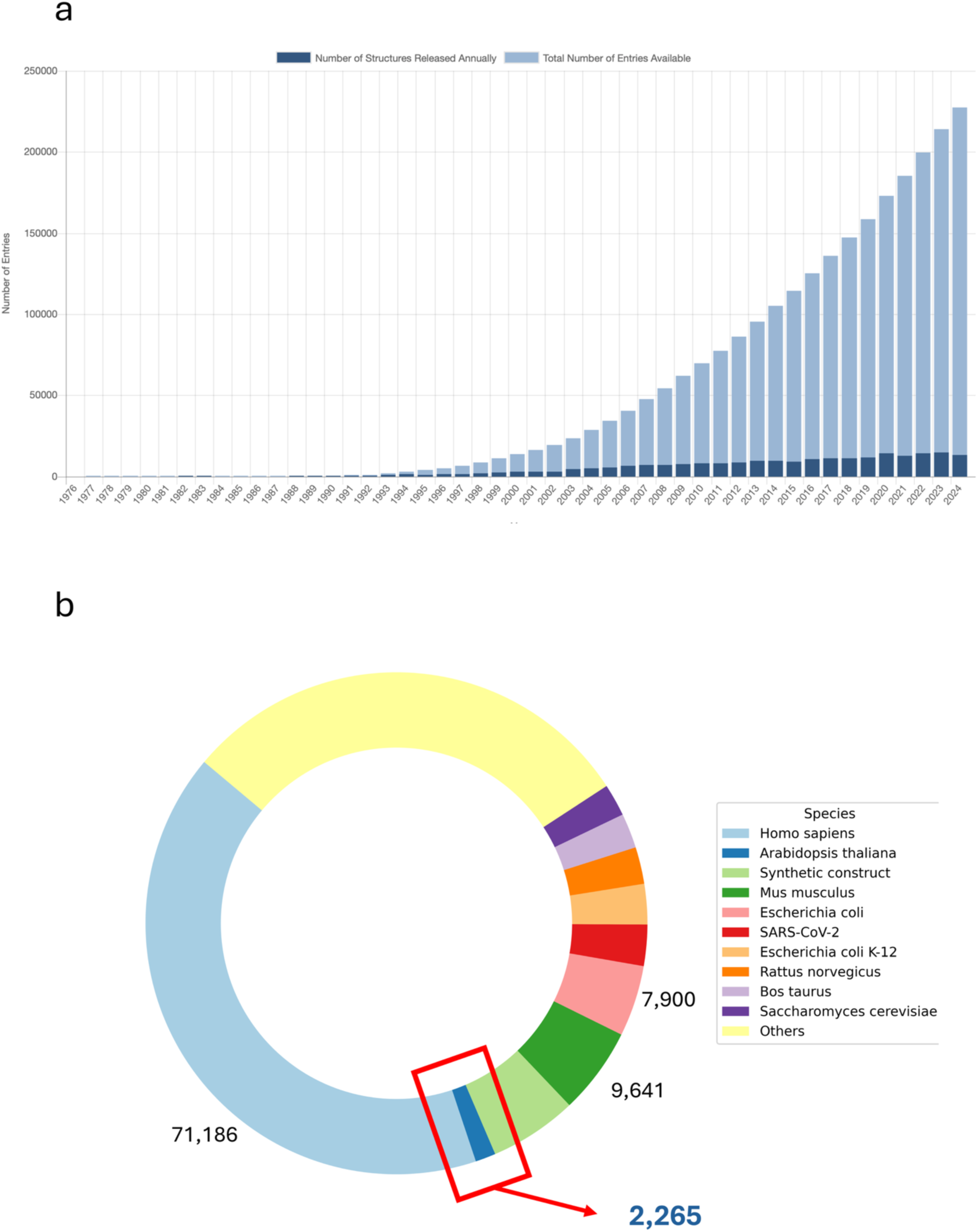
Analysis of structural coverage in Protein Data Bank (PDB) **(a**) Year-wise growth in the number of experimentally determined protein structures deposited in the Protein Data Bank (Source PDB) (b) Species-wise distribution of structures in the PDB.

To systematically quantify this gap, we analyzed 16,389 reviewed *A. thaliana* UniProt entries (from a total of 39,728 reviewed and unreviewed entries (Supplementary file 1). Of these, only 1,038 proteins (6.33%) have experimentally resolved structures in the PDB (Supplementary file 2), indicating that a large proportion of the proteome lacks structural characterization. Focusing specifically on developmental processes, we used PlantGSEA—a gene set enrichment tool for plant biology—to map biological process annotations (Figure 3a). PlantGSEA is a plant gene set enrichment analysis tool to identify the biological meaning of genes using various previously known gene sets [15]. This analysis identified 3,485 proteins associated with plant developmental processes (Supplementary file 3), of which only 337 (9.67%) have corresponding PDB structures (Supplementary file 4). These findings clearly show that experimentally resolved structures cover only a small subset of functionally critical plant proteins (Figure 3b).

**Figure 3:**
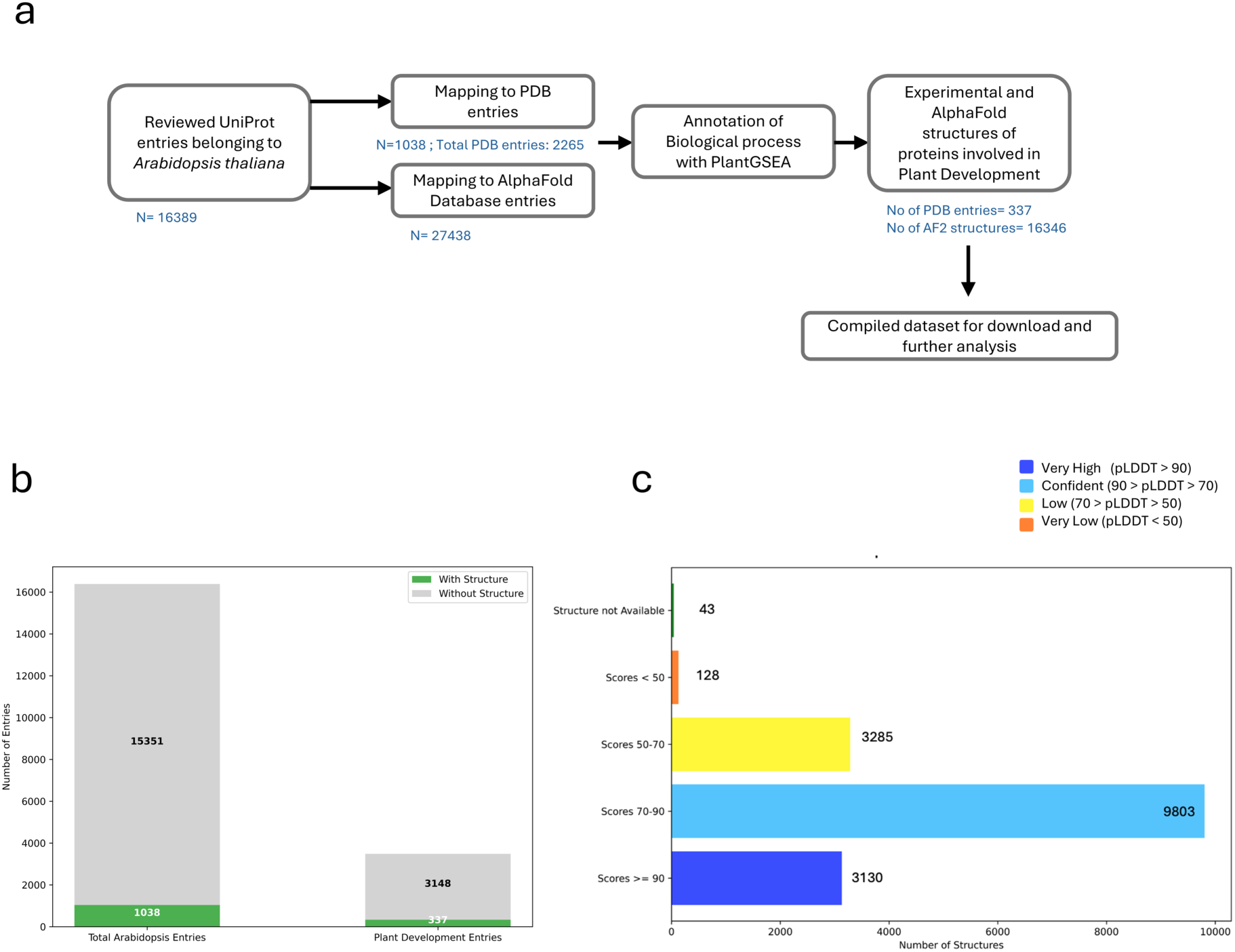
Data analysis and reviewing the structural coverage across PDB and AlphaFold (a) Workflow used for data extraction and analysis of structural coverage, (b) Number of *Arabidopsis thaliana* proteins with and without experimentally determined structures, along with the subset of plant development-related proteins and their structural availability, (c) Distribution of confidence scores (pLDDT) for AlphaFold predicted protein structures of *Arabidopsis thaliana*.

We next examined structural predictions from the AlphaFold database. Of the 39278 total *Arabidopsis thaliana* UniProt entries (including both reviewed and unreviewed), 27437 proteins have AlphaFold-predicted structures, corresponding to an overall structural coverage of ∼69.85%. Notably, this level of coverage exceeds that reported for other model organisms such as *Homo sapiens* and *Mus musculus* (Table 1). Among the 16,389 reviewed UniProt entries for *Arabidopsis thaliana*, 16,346 proteins (99.7%) have AlphaFold-predicted models. Furthermore, all 3,485 proteins associated with plant development are covered (100%) by AlphaFold predictions (Supplementary file 5). To evaluate the reliability of these predictions, we analyzed the confidence scores (pLDDT) for these reviewed entries. Approximately 79.15% of the predicted structures exhibit confidence scores above 70%, indicating generally high model confidence (Figure 3c). Collectively, these findings highlight a substantial shift between the pre- and post-AlphaFold eras of structural biology. Prior to AlphaFold, only a limited number of *Arabidopsis thaliana* proteins had experimentally determined structures, especially those involved in plant development. This limited our understanding of their mechanisms. The advent of AlphaFold has dramatically expanded structural coverage, enabling large-scale structural and functional analyses. In the next section, we discuss how experimental structural studies in the pre-AlphaFold era helped in understanding plant protein functions.

**Table 1.** Structural coverage across different organisms in the AlphaFold Protein Structure Database*.

| Species | Common Name | Reference Proteome | Total entries in UniProt | AlphaFold Predicted Structures |
| --- | --- | --- | --- | --- |
| *Arabidopsis thaliana* | *Arabidopsis* | UP000006548 | 39,278 | 27,434 |
| *Candida albicans* | *C. albicans* | UP000000559 | 6,036 | 5,974 |
| *Danio rerio* | Zebrafish | UP000000437 | 46,554 | 24,664 |
| *Dictyostelium discoideum* | Dictyostelium | UP000002195 | 12,726 | 12,622 |
| *Drosophila melanogaster* | Fruit fly | UP000000803 | 22,032 | 13,458 |
| *Escherichia coli* | *E. coli* | UP000000625 | 4,401 | 4,363 |
| *Glycine max* | Soybean | UP000008827 | 74,861 | 55,799 |
| *Homo sapiens* | Human | UP000005640 | 83,413 | 23,391 |
| *Methanocaldococcus jannaschi* | *M. jannaschii* | UP000000805 | 1,787 | 1,773 |
| *Mus musculus* | Mouse | UP000000589 | 54,747 | 21,615 |
| *Oryza sativa* | Asian rice | UP000059680 | 48,897 | 43,649 |
| *Rattus norvegicus* | Rat | UP000002494 | 47,921 | 21,270 |
| *Saccharomyces cerevisiae* | Budding yeast | UP000002311 | 6,060 | 6,039 |
| *Schizosaccharomyces pombe* | Fission yeast | UP000002485 | 5,279 | 5,128 |

\*Source: AlphaFold Database

### Experimental structural studies of *Arabidopsis thaliana* plant developmental protein for unraveling plant mechanisms – “The Pre-AlphaFold era”

Plant developmental proteins can be categorized based on their functional roles in physiological processes. Here, we highlight key advancements in structural studies of *Arabidopsis thaliana,* with particular emphasis on experimentally resolved structures. These studies demonstrate how structural biology provides critical insights into the molecular architecture of plant developmental proteins and their roles in regulating physiological processes.

The timely transition between plant developmental stages, such as the vegetative and flowering stages, is tightly regulated and is critical for reproductive success. This transition is governed at the transcriptional level through histone modification recognition by EARLY BOLTING IN SHORT DAY (EBS) [16]. It functions as a negative transcription regulator that prevents early flowering by forming a complex with H3K27me3 and H3K4me3 [17]. The binding of EBS bivalent bromo-adjacent homology (BAH) and plant homeodomain (PHD) reader modules to the H3K27me3 and H3K4me3, respectively, is crucial for the prevention of early flowering. The crystal structure of EBS domains in complex with H3K27me3 (PDB: 5Z8L) (UniProt ID: F4JL28) and EBS-H3K4me3 (PDB: 5Z8N) (UniProt ID: F4JL28) peptides provides structural insights into the chromatin-associated switch of repressive and activating signals, precisely regulating gene expression to enable timely flowering (Figure 4a). It also indicates that the binding preference of EBS by autoinhibition mode to the two antagonistic histone modifications determine the developmental outcome [16]. Notably, the AlphaFold-predicted structure of the EBS domain (AF-F4JL28 -F1) shows high confidence (pLDDT ∼86), supporting its reliability in capturing domain organization and fold (Figure 4a). Together, these findings underscore the significance of transcription regulation and post-translational histone modifications in developmental transitions.

**Figure 4:**
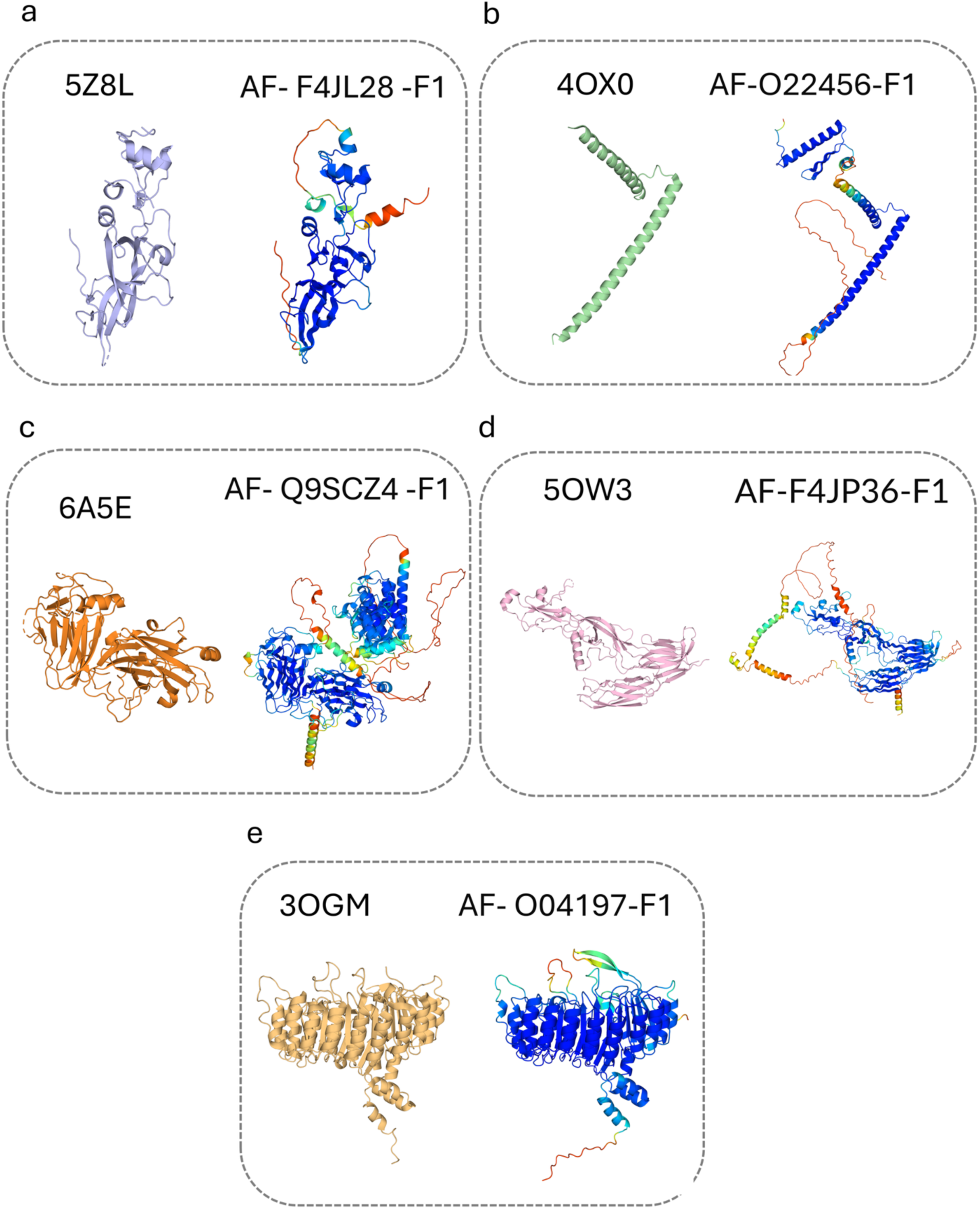
Comparison of experimentally determined crystal structures (left) and AlphaFold predicted models (right) (a) EARLY BOLTING IN SHORT DAY complex (EBS), (b) SEPALLATA3, (c) FER-LLG2-RALF23 Complex, (d) HAPLESS 2 (HAP2), (e) Coronatine-insensitive protein1 (COI 1). Note: Predicted structures are colored by pLDDT confidence scores. Corresponding PDB IDs and AlphaFold entries are labeled.

Plant gene expression during development is further controlled by MADS-domain transcription factors, which form a complex regulatory network [18]. These proteins have undergone extensive diversification in plants through gene duplication and domain variation. Members of the MADS-box family regulate distinct gene sets by forming dimers and tetramers, orchestrating floral organ specification, including sepals, petals, stamens, carpels, and ovules. [19]. The highly variable binding specificity and selectivity of Class E MADS-box domain proteins, such as the central floral regulatory protein SEPALLATA3 (SEP3), have been structurally studied by Puranik et al. in 2014, elucidating the 2.5-Å crystal structure (PDB: 4OX0) (UniProt ID: O22456) of its K domain and some portion of its I domain in *Arabidopsis* [20] (Figure 4b). The study identifies two amphipathic helices connected by a kink, with one helix promoting dimer formation and the other involved in tetramer formation. Hydrophobic residues critical for oligomerization and DNA binding were also identified, highlighting the structural basis of transcriptional regulation. The comparative AlphaFold structure of SEPALLATA3 (AF-O22456-F1) shows a similar folding pattern to the experimentally determined structure, with a high confidence score of 76.31%, providing reliability of AlphaFold for structure prediction (Figure 4b). These findings provide a mechanistic framework for understanding floral development and offer avenues for targeted manipulation of flower traits in crop improvement.

Pollination in angiosperms is a tightly regulated process, in which pollen acceptance and rejection responses ensure species integrity and genetic diversity. To prevent the germination and hydration of incompatible pollen, members of the CrRLK1L subfamily of receptor kinases – FERONIA (FER1), along with GPI-anchored co-receptor LORELEI-LIKE proteins (LLG1/LLG2), and stigmatic cationic peptides Rapid Alkalinization Factor (RALF) form a heterotrimeric complex [21]. A study by Xiao et al. elucidated the mechanism by which the stigmatic peptide RALF23 binds to the FER1-LLG1/2 complex, triggering an increase in reactive oxygen species (ROS) in the stigma and inhibiting pollen attachment and hydration [22]. The crystal structure of RALF23 in complex with FER1 and LLG1 (PDB: 6A5E) (UniProt ID: Q9SCZ4) (Figure 4c) has revealed that the negatively charged residues of the FER1-LLG1 complex interact with the N-terminal region of RALF23 to initiate the signal of Reactive Oxygen Species (ROS) production via membrane-localized Respiratory burst oxidase homologs (RBOHD/F) NADPH oxidase. The extracellular domain of FER1 displays a tandem malectin-like domain conformation, forming a scaffold that lacks a canonical peptide-binding pocket, which, due to its rigidity, hinders the direct binding of RALF23 to FER1 alone. In contrast, the LLG1 structure features a compact α/β fold with a surface groove that serves as a docking site for RALF23. The crystal structures show that RALF23 binds in an extended yet ordered conformation, with its N-terminal residues deeply embedded within the LLG1 groove, stabilized by extensive hydrogen bonds, salt bridges, and hydrophobic contacts. The C-terminal region of RALF23 remains more flexible but contributes to complex stabilization by reinforcing LLG1–FER1 interactions. Importantly, FER1 engages the LLG1–RALF23 module through defined contact surfaces on its malectin-like domains, resulting in a heterotrimeric assembly in which the peptide acts as a molecular attachment rather than a classical receptor ligand. The structural comparisons further revealed conformational plasticity in LLG1 loops, providing a basis for recognition of the RALF peptide. These crystallographic studies established a mechanistic model in which ligand-induced co-receptor engagement and receptor assembly regulate FER1 signaling during *Arabidopsis* development rather than direct ligand binding to the kinase receptor [22]. This study has elucidated the structure of the complex of the extracellular domain of FER1 with the molecular partners and its binding mechanism in self-incompatibility. The structure of the full-length FER1 protein along the cytosolic domain is yet to be elucidated, which may provide critical insights into downstream signaling pathways. The AlphaFold structure of FER1 (AF- Q9SCZ4 -F1), as displayed in Figure 4c, has a multidomain structure with a high pLDDT score 79.81%, whereas the experimental structure of FER1 represented only the extracellular domain involved in the self-incompatibility (SI) mechanism. These results highlight the utility of computational approaches in addressing challenges associated with structurally complex, multidomain proteins.

Understanding the mechanisms of male and female gamete fusion is fundamental to studying sexual reproduction in plants. HAPLESS 2 (HAP2) is a crucial protein from the male gamete involved in interacting with the female gamete membrane. The X-ray crystallographic study of HAPLESS2 (HAP2) from *Arabidopsis thaliana* by Fedry et al. uncovered the previously unclear mechanism of HAP2 and female gamete membrane fusion [23]. The C-terminally truncated crystal structure of the trimeric ectodomain (PDB: 5OW3) (UniProt ID: F4JP36) (Figure 4d) revealed an overall fold resembling that of viral fusion proteins. However, unlike viral counterparts, HAP2 mediates membrane fusion through an amphipathic helix that inserts into the female gamete membrane. In contrast, HAP2 proteins in trypanosomes utilize multiple loops for membrane interaction, underscoring mechanistic divergence across taxa despite structural similarity. The AlphaFold-predicted structure of HAP2 (AF-F4JP36-F1) exhibits a high confidence score (pLDDT = 76.94%), further supporting AlphaFold’s accuracy and reliability in structure predictions (Figure 4d).

Complex biological processes in plants are orchestrated through highly coordinated interactions among proteins and other biomolecules. Proteins expressed at specific developmental stages assemble into functional complexes, generating synergistic effects that drive precise physiological outcomes [5]. Structural characterization of such complexes provides critical insights into molecular assembly, binding specificity, and regulatory mechanisms, and has enabled the development of innovative strategies to address biological challenges. Jasmonic acid, a key phytohormone, regulates plant growth, development, and defense responses. Perception of jasmonate signals involves a co-receptor complex comprising the CORONATINE INSENSITIVE 1 (COI1) F-box protein, JASMONATE ZIM DOMAIN (JAZ) transcriptional repressors, and small-molecule ligands [24]. Structural analysis by Sheard et al. revealed a unique mechanism for the perception of bioactive jasmonoyl-L-isoleucine (JA-Ile) perception by the COI1–JAZ co-receptor complex [25]. The 3.34 Å crystal structure (PDB: 3OGM) (Uniprot: O04197) (Figure 4e) showed that the COI1–ASK1 complex forms a ligand-binding pocket that accommodates the bipartite degron of JAZ. A conserved α-helix of JAZ docks into this binding pocket, effectively trapping the JA-Ile hormone. Additionally, it has been recently discovered that inositol pentakisphosphate (IP₅) cooperatively binds, acting as a cofactor that promotes complex formation and stabilization. The AlphaFold-predicted structure of COI1 (AF- Q39255 -F1) exhibits a high confidence score (pLDDT ≈ 91.6%), further demonstrating the accuracy of structure prediction approaches (Figure 4e). This study underscores the importance of structural analyses in unraveling the complexity and functional dynamics of hormone signaling pathways.

Here, we have discussed structural studies of a few representative plant developmental proteins that have provided insights into their functional roles in physiological processes. The crystallographic study of a monomeric protein and the study of functional domains provide insights into active sites, biochemical mechanisms, and the developmental processes associated with them. The protein complexes with co-factors or other binding protein partners shed light on the molecular assembly pattern, binding affinities, and functional mechanisms of the pathway (Figure 1). Nevertheless, many proteins lack full-length experimentally resolved structures due to intrinsic disorder, conformational flexibility, or the presence of hydrophobic transmembrane regions. Addressing these challenges will require advances in experimental techniques such as cryo-electron microscopy, complemented by *in silico* approaches. (Figure 1). The integration of experimental and computational methods will be essential for a comprehensive understanding of plant biological systems and for guiding strategies to enhance crop productivity and stress resilience.

### Advancements in structural studies of *Arabidopsis thaliana* plant developmental proteins-insights from “The Post-AlphaFold era.”

Recent advances in experimental approaches for resolving protein structures have significantly improved our understanding of protein behavior *in vivo*. However, these methods are often labor-intensive and demand sophisticated instrumentation, technical expertise, and substantial time investment. Computational approaches for predicting protein structures provide an efficient, and systematic alternative, reducing reliance on time-consuming experimental workflows [26, 27]. The conventional computational approach to predict protein structure using homology modeling requires significant sequence similarity to already elucidated protein structures [28]. Hence, the structure prediction of a protein lacking any sequence identity to known protein structures has been difficult. In contrast, ab initio methods and advanced AI-based tools, such as AlphaFold 3 and its predecessors, enable accurate prediction of a protein’s three-dimensional structure even in the absence of homologous templates [9, 29]. Here, we compare AlphaFold-predicted structures of protein lacking experimentally determined crystal structures to evaluate the accuracy and reliability of this approach and its contribution to understanding protein function and interactions.

A critical prerequisite for successful fertilization is the formation of viable pollen. Early post-meiotic pollen development and maturation depend on coordinated tapetal development, followed by cytosolic and cell wall remodeling within pollen grains. A key regulator of these processes is the transcription factor MS1 (Male Sterility 1), which ensures proper pollen maturation and fertility (Figure 5a) [30–35]. It’s an integral protein that regulates one of the key early steps in plant reproductive development and ensures fertility. Although the functional relevance is known, the experimental structural information remains limited. The *in silico-*predicted structure by AlphaFold provides high confidence and pLDDT scores for the 672-amino-acid-long protein (Figure 5a) (Uniprot ID: Q9FMS5). The structure reveals a characteristic PHD-Finger domain at the C-terminus and a central helix-turn-helix domain, indicating its importance as a DNA-binding transcriptional activator. Understanding the structural features of such proteins would provide more insights into their binding dynamics, interaction partners and functional relationships.

**Figure 5:**
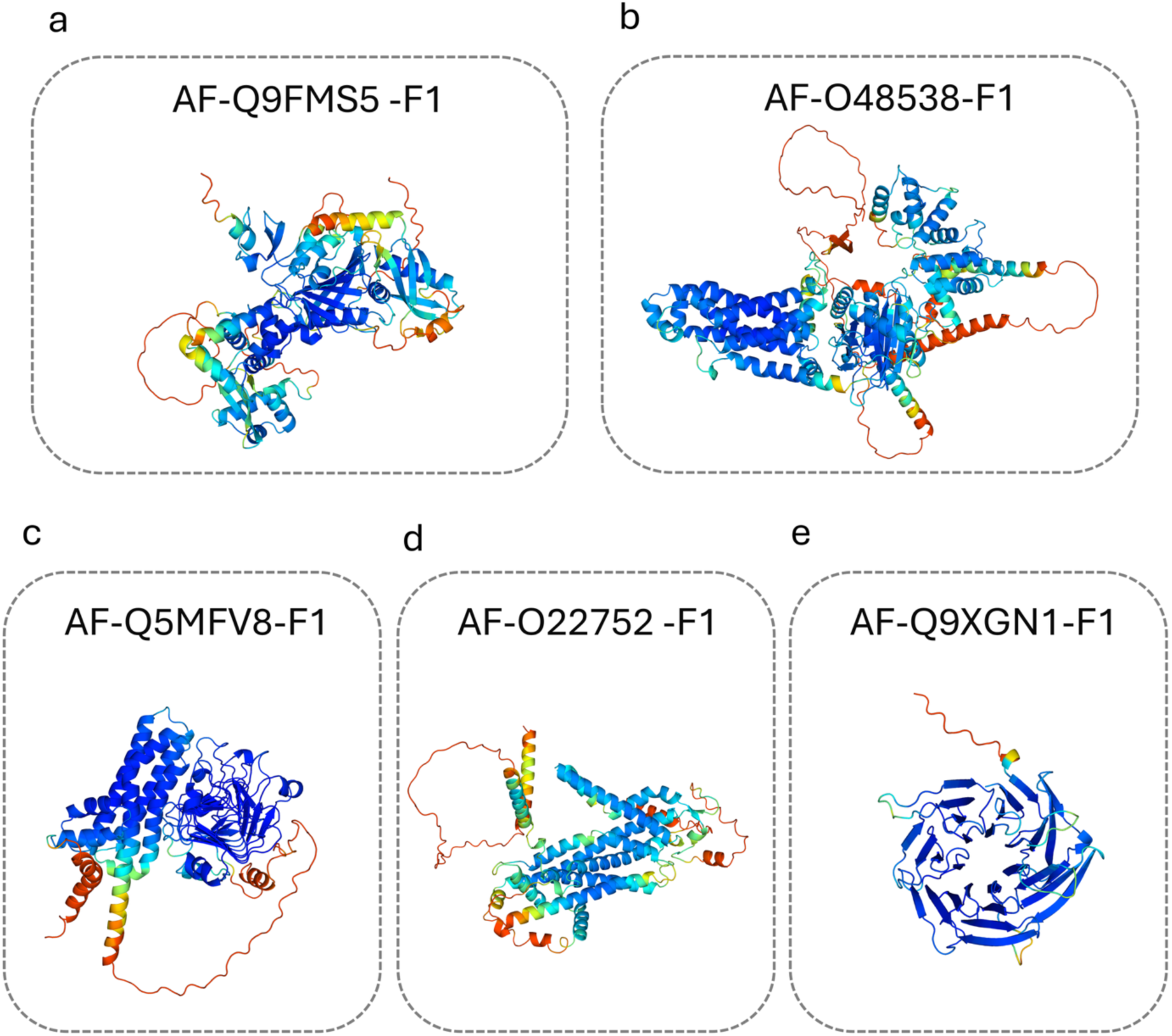
Representation of AlphaFold predicted structures for proteins without experimentally determined structures (a) MS1, (b) RBOHD/F, (c) PME 5, (d) Nortia, (e) TTG1. Note: Models are colored by pLDDT confidence scores, with corresponding AlphaFold IDs provided.

Various plant species, including *Arabidopsis,* have evolved mechanisms to maintain genetic diversity, promote outcrossing, and avoid inbreeding depression [36, 37]. One of the integral mechanisms is SI; although lost in *A. thaliana,* it persists in various *Arabidopsis* specie*s* [38, 39]. ROS act as key regulators of SI, modulating pollen acceptance or rejection through control of pollen hydration and attachment [40, 41]. The pathway is mediated by FER, which activates downstream signaling involving RBOHD/F [40, 41], an NADPH oxidase that generates superoxide and leads to pollen rejection [37]. Although an integral regulator of self-incompatibility, studied functionally over the past decade, RBOHD/F has not been crystallized, lacking to provide insights into its structural regulation. AlphaFold predictions of this 944-amino-acid protein exhibit high confidence (Figure 5b) (Uniprot ID: O48538). The structure possesses varying subsequent cytoplasmic, transmembrane, and extracellular regions. The N-terminal EF-hand domains, which mediate Ca²⁺ binding and are critical for ROS regulation [42], are predicted with lower confidence, likely reflecting their intrinsic flexibility. In contrast, the C-terminal catalytic regions, including the ferric oxidoreductase and FAD-binding domains essential for ROS production [43], are predicted with very high confidence (Figure 5b). These structural insights, although computational, provide a valuable basis for understanding SI regulation and may inform strategies for hybrid breeding and crop improvement [44]. Beyond SI, RBOHD/F also plays broader roles in ROS-mediated signaling during hypersensitive responses, stomatal closure, and responses to UV-B radiation and abscisic acid [45–49].

Another crucial aspect of plant reproduction is the precise delivery of sperm cells to the egg. Following pollen deposition, compatible pollen–stigma interactions initiate adhesion, hydration, and germination. The pollen tube then penetrates the stigma and style, navigates the transmitting tract, and ultimately reaches the ovule, making this a highly coordinated and complex process [50–59]. Pectin methylesterases (PMEs) expressed in pollen and pollen tubes are key regulators of this process, modulating cell wall properties to facilitate tube growth within female tissues [60]. Given the intrinsic nature previously studied for these PMEs [60], understanding their structural mechanisms and dynamics, as well as their structure-function relationships, is integral to manipulating reproductive success. Two decades have passed since the functional attributes were identified, though less progress has been made in resolving their crystal structure. *In silico* structure prediction powered by AlphaFold has predicted the structure with high confidence and pLDDT scores (Figure 5c) (UniProt ID: Q5MFV8). The 595 amino acid-long protein has been predicted, with most regions assigned very high confidence, and some portions of the N-terminal, mostly the signal peptide, assigned low to very low confidence, as indicated by the pLDDT scores. Based on previous studies, the N-terminal region, including the pectin methyl esterase inhibitor (PMEI) domain, was predicted with high confidence to regulate untimely PME activity, acting as an autoinhibitory domain [61]. This domain is targeted by subtilisin-like protease (S1P) cleavage prior to cell wall targeting [61]. Further, the C-terminal pectinesterase domain, also predicted with high confidence, mediates pectin demethylesterification, a process essential for pollen tube growth and guidance (Figure 5c) [60]. Although the functions and regulations have been previously known, the introduction of the predicted structure highlights the key attributes and the topology of the integral domains necessary for manipulating reproductive success and crop yield.

Pollen tube, upon exiting the transmitting tract, is attracted to the ovules in a one-on-one fashion. The subsequent arrival at the embryo sac and the release of two sperm for double fertilization lead to zygote formation and endosperm development. One of the players integral to the process is NORTIA (NTA), which, alongside FER, regulates pollen tube reception [62]. Both genes have been implicated in pollen tube reception and pathogen resistance [62]. Although it is integral for successful reproduction and double fertilization, the experimental structural attributes remain limited. AlphaFold has predicted the structure of NORTIA, a member of the MLO (Mildew Resistance Locus O) family, which comprises seven transmembrane domains, a characteristic of G-Protein Coupled Receptors (GPCRs) (Figure 5d) (Uniprot ID: O22752) [62, 63]. The 542-amino-acid structure was predicted with high confidence, indicating that the calmodulin-binding domain is located toward the C-terminus. The domain has previously been validated for its role in binding to calmodulin, leading to recruitment to the filiform apparatus, regulation of Ca^2+^ channel activity, and pollen tube reception (Figure 5d) [64–67]. Given its integral role in pollen tube reception and pathogen defense, *in silico* structure prediction provides impetus for further understanding the structural attributes of this molecular control and provides evidence for its evolution [62].

Post-fertilization, seed development ensures the propagation of the next generation through the accumulation of storage reserves and protective structures. One of the key players regulating seed coat mucilage production and the accumulation of fatty acids and proteins is Transparent Testa Glabra 1 (TTG1) [68–70]. This gene is also known to regulate trichome and root hair formation, flavonoid biosynthesis, flowering time, and responses to biotic and abiotic stress [71–78]. Given its integral role across various developmental aspects of plants, it is important to understand its structural attributes. Although no experimental structure is currently available, AlphaFold predictions of the 341-amino-acid TTG1 protein (UniProt: Q9XGN1) show high confidence and reveal characteristic WD-repeat motifs (Figure 5e) [73]. These repeats facilitate interactions with MYB and bHLH transcription factors to form the MYB–bHLH–WD40 (MBW) complex, a key regulatory module in plant development (Figure 5e) [79–83]. Given the importance and roles of this protein, insights into its structural modalities provide a better understanding of its interactions and functional relevance.

Collectively, these examples underscore the importance of understanding protein structure–function relationships in plant biology. While experimentally determined structures have provided invaluable insights over past decades, many proteins remain structurally uncharacterized due to technical limitations. The emergence of advanced *in silico* tools, particularly AlphaFold, has transformed this landscape by enabling high-confidence predictions of complex protein structures (Figure 1). These models not only reveal structural features but also illuminate molecular mechanisms, interaction interfaces, and functional dynamics, highlighting the growing significance of integrative structural biology approaches in addressing fundamental and applied challenges in plant science.

### Challenges in the structural study of plant developmental proteins and future directions

Researchers face several challenges in the structural study of plant developmental proteins, primarily due to the structural complexity, dynamics, and methodological constraints that limit progress. Proteomics studies have previously identified a wide array of plant proteins involved in various developmental stages [84]. The expression and concurrent localization of certain proteins in different plant parts during plant development suggest diverse functionality; however, this complicates the understanding of their complex physiological roles in biological processes [85]. Hence, it is a prerequisite to study protein structure in both the monomeric state and in complexes to understand their physiological roles. The primary challenge in experimentally studying plant protein structures is their low solubility, folding patterns, protein yields, and PTMs [86]. The PTMs of plant proteins play a pivotal role in protein localization within different compartments, enzymatic activity, functionality, and interactions with other biomolecules, thereby regulating physiological processes [87]. They also regulate protein fate, turnover rates, and molecular conformations [88]. Furthermore, they pose significant challenges for heterologous protein expression and purification, including the formation of unstable, misfolded, or inactive protein forms and heterogeneity in PTMs when expressed in unsuitable expression systems such as prokaryotes and yeasts [89]. Employing the baculovirus-mediated insect cell line expression system and *in planta* transient expression for plant protein production offers high accuracy in eukaryotic PTMs and a safe alternative with higher production rates [90–92]. Intrinsically disordered regions in plant proteins are functionally active regions lacking ordered or defined three-dimensional structures, yet performing their physiological roles [93]. Although these regions play a critical role in modulating the binding affinities, structural flexibility, and functional versatility of plant proteins, they pose significant challenges for structural studies. Transcription factors in plant systems are well known for harboring functional intrinsically disordered regions, which are imperative for dynamic interactions with other biomolecules and for regulating the transcriptional machinery [94]. These proteins do not fit the standard definition of proteins suitable for crystallographic or NMR studies, i.e., the native conformation of a protein with a spatial arrangement of amino acid residues that provides a specific structure to the active region and a suitable microenvironment for enzyme activity [95]. There is a need for efficient expression and purification methods for proteins with intrinsically disordered regions, as they are more prone to proteolytic degradation and the formation of insoluble aggregates, resulting in reduced yields. The combined analysis of intrinsically disordered proteins using high-resolution NMR and cryo-EM data can address structural challenges [96]. The developmental membrane proteins are usually membrane-localized or membrane-anchored multidomain proteins acting as receptor molecules and signal transducers [97]. They play a crucial role in cell differentiation, plant growth, reproduction, and environmental responses [98]. The structural study of plant membrane proteins with full-length structures is difficult to resolve due to the hydrophobic membrane-bound regions, which lack *in vitro* solubility and stability. The flexible domains of the protein devoid of a static conformation exhibit conformational heterogeneity in solution, making it difficult to crystallize [99]. Plant-specific PTM patterns in membrane proteins, such as glycosylation, SUMOylation, and phosphorylation, may differ in heterologous systems compared to in-planta conditions [100]. Given the challenges associated with full-length plant membrane protein structures, there is an urgent need to develop effective solutions through combined experimental and *in silico* strategies. Strategies such as membrane mimics and nanodiscs are considered advantageous for enhancing the stability and functional activity of membrane-associated protein structures, which may be helpful for studying plant proteins structurally [101]. The cryo-EM technique is emerging as a more reliable method for studying oligomeric, dynamic membrane protein structures that are usually non-crystallizable. Unlike X-ray crystallography, proteins can be flash-frozen by vitrification and multiple orientations, and conformational states can be captured to analyze data and to identify functionally active forms, along with liposome- or nanodisc-associated complexes [102, 103]. To deduce the structural information of proteins, it is crucial to advance AI-driven methods and to explore bioinformatic tools such as AlphaFold to address the challenges posed by intrinsically disordered proteins, membrane proteins, complexes, and difficult-to-crystallize proteins [104] The advent of AlphaFold and the advancement in various AI-based prediction tools have significantly enhanced the coverage and confidence in predicted protein models. Furthermore, the “protein folding problem” – predicting the 3D shape of proteins from one-dimensional amino acid sequence, has certainly been resolved through the advancements in the last decade, with emphasis on AI-based *in silico* tools. The ease of prediction, aligned with more confident structures have drastically increased the utility and availability of these tools. Almost three decades of continuous advancement and improvement have led to tools such as AlphaFold, ROSETTA, and others that provide significant support for structure prediction, interactions, and functional relevance. Furthermore, a shift from traditional modeling to AI-based tools has been observed over the last decade in the CASP (Critical Assessment of Protein Structure Prediction). These tools are essentially upgrades and aid to the expensive, time-consuming experimental methods, addressing their limitations. Utilizing these tools will help in enhancing structural coverage, understanding, and elucidating key phenomena and physiological processes.

## Conclusion

The structural and functional characterization of plant developmental proteins is crucial for developing innovative strategies to address critical challenges in crop improvement and agricultural sustainability. A marked disparity between the availability of experimental structural data for *Arabidopsis thaliana* and *Homo sapiens* proteins highlights the pressing need to prioritize plant protein structural biology. Systematic mapping of UniProt entries to the Protein Data Bank (PDB) for *Arabidopsis thaliana* developmental proteins reveals a substantial gap in structural coverage. This limited coverage reflects inherent challenges, including the large size and multimeric nature of many plant proteins, complex molecular assemblies, diverse post-translational modifications (PTMs), low solubility, structural dynamics, membrane association, and the prevalence of intrinsically disordered regions. Addressing these challenges will require innovative experimental strategies, such as baculovirus-mediated expression systems, advanced purification techniques, and the use of nanodiscs or membrane mimetics, alongside high-throughput structural approaches including X-ray crystallography, nuclear magnetic resonance (NMR), and cryo-electron microscopy (cryo-EM). Our analysis indicates that only 6.33% of *Arabidopsis thaliana* proteins represented in UniProt have experimentally resolved structures, whereas AlphaFold provides structural predictions for approximately 99.7% of entries. Similarly, proteins specifically involved in plant development show 9.67% experimental structural coverage compared to nearly complete (100%) coverage through AlphaFold predictions. Notably, the breadth and confidence of AlphaFold predictions for *Arabidopsis thaliana* proteins surpass those reported for *Homo sapiens*, underscoring its transformative potential in plant structural biology. Hence, integrating experimental and computational methods will pave the way for overcoming the challenges in plant protein structural biology. Nonetheless, our study provides a comprehensive overview of the structural coverage of *Arabidopsis thaliana* plant developmental proteins, highlighting differences in coverage between the pre- and post-AlphaFold eras. The datasets and analyses presented here offer valuable resources for exploring structure–function relationships and guiding future research. Importantly, they emphasize the need for continued methodological advancements—both experimental and *in silico* to meet the growing demand for detailed structural insights and to translate these findings into practical applications for crop improvement.

## Statement and Declarations

### Funding

This work was supported by the Department of Biotechnology (DBT) RA fellowship to SSR, IIT Gandhinagar excellence fellowship to KS, MoE Prime Minister Research Fellowship to HB, DBT-Ramalingaswami fellowship to AS and SS, SERB-Start up Research Grant, and a start-up grant from the Indian Institute of Technology Gandhinagar to SS.

## Acknowledgments

We acknowledge the Department of Biotechnology Research Associate for the fellowship to SSR, IITGN for the fellowship to KS, and the Ministry of Education for the Prime Minister Research Fellowship to HB. We acknowledge support from DBT for the Ramalingaswami Re-entry fellowship and IITGN start-up grants to AS and SS.

## Author contributions

SS conceived and designed the outline, supervised, edited, and proofread the manuscript. AS conceived, supervised, and proofread the manuscript. SSR, KS, and HB wrote the first draft of the manuscript. All the authors were involved in discussions and commented on the manuscript.

## Conflicts of interest

The authors declare no conflict of interest.

## Availability of data and materials

The datasets supporting the conclusions of this article are included within the article and its supplementary files.

## Notes

### Competing Interest Statement

The authors have declared no competing interest.

